# Integrating diverse statistical methods to analyse stage-discriminatory cell interactions in colorectal neoplasia

**DOI:** 10.1101/2024.06.02.597010

**Authors:** Joshua A. Bull, Eoghan J. Mulholland, Joshua W. Moore, Jesús J. Bosque, Bernadette J. Stolz, Joseph Boen, Holly R. Eggington, Hayley L. Belnoue-Davis, Helen Jones, Chandler D. Gatenbee, Alexander R. A. Anderson, Alistair Easton, Peter Todd, Christopher Cunningham, Stephen Taylor, Helen M. Byrne, Simon Leedham

**Affiliations:** Wolfson Centre for Mathematical Biology, Mathematical Institute, University of Oxford, Oxford OX2 6GG, UK; Centre for Human Genetics, Roosevelt Drive, University of Oxford, Oxford, UK; Department of Radiation Oncology, Massachusetts General Hospital and Harvard Medical School, Boston, MA, USA; Laboratory for Topology and Neuroscience, EPFL, Station 8, 1015 Lausanne, Switzerland; Nuffield Department of Surgical Sciences, University of Oxford, UK; Department of Integrated Mathematical Oncology, H. Lee Moffitt Cancer Center & Research Institute, 12902 Magnolia Drive, SRB 4, Tampa, FL, 336122, USA; Department of Oncology, Old Road Campus Research Building, Roosevelt Drive, University of Oxford, Oxford, UK; Ludwig Institute for Cancer Research, Nuffield Department of Medicine, University of Oxford, Oxford OX3 7DQ, UK; Translational Gastroenterology Unit, John Radcliffe Hospital, University of Oxford, and Oxford NIHR Biomedical Research Centre, Oxford, UK

**Author notes:** These authors contributed equally to this work (Joshua A. Bull, Eoghan J. Mulholland, Helen M. Byrne, Simon Leedham). Contact.

**Keywords:** Spatial biology, colorectal neoplasia, multiplex imaging, spatial statistics, topological data analysis, spatial analysis

## Abstract

Spatial biology has the potential to unlock information about the disrupted cellular ecosystems that define human disease. Quantitative analysis of spatially-resolved cell interactions allows mapping of tissue self-organisation and assessment of why cells interact differently in physiological and pathological contexts. However, the complexity of mammalian tissues, that occur across a spectrum of length scales, presents significant challenges for spatial analysis, increasing the gap between our capacity to generate and biologically interpret these datasets. Here, we have adapted a range of mathematical tools to develop a suite of spatial descriptors, and deployed them to determine how cell interactions change as colorectal cancer progresses from benign precursors. We demonstrate that combining mathematical analyses permits insightful examination of tissue organisational structures and identifies variable cell-interaction pathways that underpin disease progression. Mathematical tool triangulation can cross-corroborate spatial biology findings, facilitating development of analysis pipelines that are robust to individual method limitations.

## Introduction

Mammalian tissues are complex cellular ecosystems, defined by interactions between embryologically-distinct cell populations (e.g epithelial, immune and stromal cells), that change with the onset and progression of disease (Schmitt and Greten 2021). The resultant disruption in tissue architecture characterises the recognizable pathological features of the condition (Anderson and Simon 2020). The emergence of spatial biology technologies (including multiplex proteomics and spatial transcriptomics) permits the quantification of cell associations, enabling the mapping of tissue self-organisation and the determination of how and why cells interact differently in physiological and pathological contexts. Multiplex imaging technologies produce spatially-resolved maps of tissue samples at single-cell resolution, distinguishing cells based on the functional markers they express. An increasing number of technology platforms now exist (Pourmaleki et al. 2023; Goltsev et al. 2018; Giesen et al. 2014), varying in the size of the tissue area that can be imaged and the number of expressed genes or protein markers measured. However, following image processing and cell segmentation, all platforms ultimately result in the definition of individual (*x,y*) coordinates of cell centres, providing information about cell numbers, densities and locations to delineate the tissue spatial context.

Measuring and understanding non-random cell interactions has driven the development of applied mathematical tools to quantify spatial relationships between cells with phenotypes of interest (Matejka and Fitzmaurice 2017; Anscombe 1973). These include simple distance measurements such as the mean minimum distance between pairs of those cells (Feng et al. 2023), or nearest-neighbour distributions (Hagos et al. 2022). More complex spatial statistics describe spatial relationships across a range of length scales by comparing observations of pairwise distances against a null model of spatial spread; for example, the J-function (van Lieshout and Baddeley 1996) compares the observed nearest-neighbour distribution of pairwise distances between points against the empty space function, in which test points are randomly sampled across the domain to simulate cell locations (Bull et al. 2020). The approach of comparing observed statistics against a statistical null distribution is common within spatial statistics (Baddeley, A., Rubak, E., and Turner, R. 2016), and a variety of such methods have been used to describe spatial relationships between cells in imaging data. Perhaps the most common such approaches are Ripley’s K function (Ripley 1977; Feng et al. 2023) and the pair correlation function (PCF) (Miroshnychenko et al. 2023; Bull et al. 2024), two related metrics which characterise the distribution of points separated by a range of distances *r*.

While these spatial statistics describe correlation between points across length scales, other approaches describe correlation within a fixed length scale. For example, ecological methods like the Morisita-Horn index (Hagos et al. 2022) and quadrat correlation matrix (QCM) have been used to characterise cell-cell relationships (Morueta-Holme et al. 2016; Gatenbee et al. 2022; Weeratunga et al. 2023) and the cellular composition of distinct tissue regions (Phillips et al. 2021; Schürch et al. 2020). Network approaches have also been used to examine spatial relationships between vascular and neuronal densities in the brain (Wu et al. 2022) and to quantify the numbers of interactions within a connectivity network (Weeratunga et al. 2023; 2024). Spatial architecture has also been characterised in terms of “topological features”, which quantify structures such as the numbers of connected components, loops in 2-dimensional data or voids in 3-dimensional data in a dataset. Topological Data Analysis (TDA) has been used to characterise such spatial distributions of immune cells in cancer (Vipond et al. 2021; Stolz et al. 2023), tumour vasculature (Stolz et al. 2022), and the extracellular matrix in lung cancer (Yoon et al. 2024). Several software approaches include spatial analysis methods in their analysis pipelines (Stoltzfus et al. 2020; Palla et al. 2022). SPIAT provides an R toolkit to integrate several cell co-localisation metrics (Feng et al. 2023). GraphCompass (Ali et al. 2024) and SCIMAP (Nirmal and Sorger 2024) provides graph based workflows for spatial analysis. However, no quantitative comparison of these spatial analysis methods has yet been conducted, because the software pipelines focus on one or two methods only.

While each of the methods discussed above has distinct strengths, a problem with applying them in isolation is that complex mammalian tissue ecosystems are defined by multi-level cell interactions that act across cellular compartments and a spectrum of length scales. Some interactions are proximity-based, mediated by direct physical contact or paracrine secreted signalling pathways - such as the interaction of a leukocyte with an antigen presenting cell (Labernadie et al. 2017). Other organisational structures occur at a tissue architectural level - such as the development of tertiary lymphoid structures at the invading edge of a cancer (Lin et al. 2023). It is presently unclear which statistical methods are best suited to encompass this organisational complexity - whether individual analytical tools succeed in assessing spatial associations across length scales, whether biological findings reported in quantitative studies could have been detected using alternative approaches, or whether complementary mathematical approaches could be combined to identify the same underlying biological features.

In this work, we adapt methods from a range of fields to develop and apply a suite of mathematical analytical tools to spatial biology datasets. We assess the performance of these tools in a human disease setting by analysing changing cell interactions as colorectal cancer progresses from paired benign precursor lesions. We demonstrate that key cell interactions quantifiably change between benign and malignant pathological states, and that these stage-discriminatory cell interactions can vary from patient-to-patient. Importantly we show that, although different mathematical methods are sensitive to different spatial features, tool triangulation can identify the same underlying cell interaction dynamics, and that the combinatorial application of complementary statistics enhances our ability to detect spatial patterning compared to the use of any one method alone. Our work suggests a different approach to spatial analysis: concordance between multiple quantitative spatial descriptors can reinforce the validity of biological interpretation, leading to more robust conclusions and increasing our confidence that detected key cell interactions are genuine and biologically relevant.

## Results

### Synthetic data demonstrate differential capacity of mathematical tools to describe spatial patterning

In order to demonstrate the benefits of using multiple mathematical tools to describe different types of spatial structure in tissue, we first generated four synthetic datasets that represent the variations in cell-cell interactions typical of tissue organisational structures relevant to intestinal cancer samples (Figure 1):

**Figure 1:**
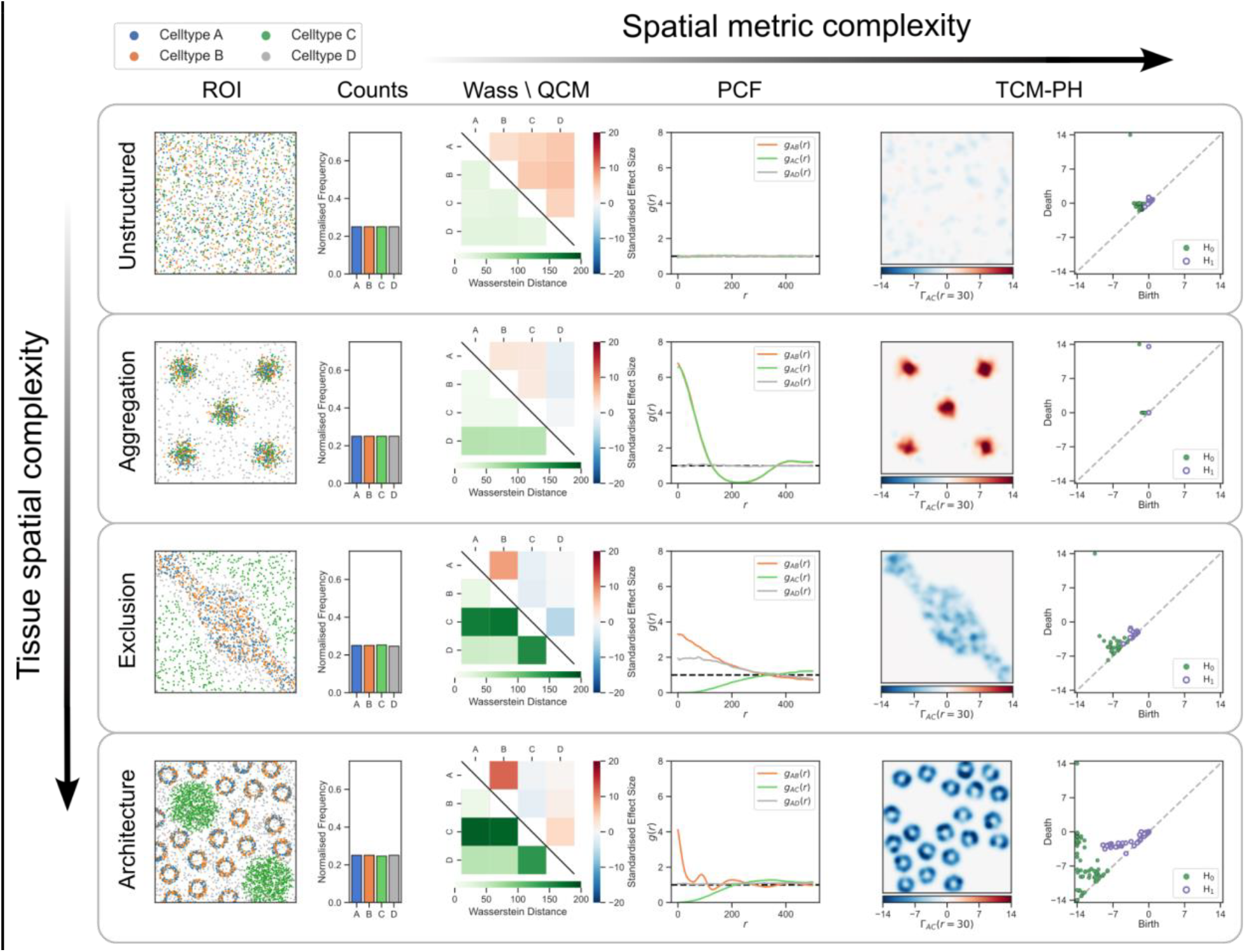
Multiple spatial metrics are needed to distinguish different spatial structures. Capturing complex spatial architecture requires a range of spatial metrics. The four synthetic datasets illustrate different spatial relationships between cell types A (blue), B (orange), C (green), and D (grey), which represent the range of cell-cell interactions in intestinal cancer tissue samples. In each ROI, type D cells are randomly distributed and the others are distributed as follows: “Unstructured,” where all cell types are randomly distributed; “Aggregation,” where cell types A, B, and C aggregate in dense clusters; “Exclusion,” where cell types A and B cluster centrally, while C is at the edges; “Architecture,” where cell types A and B form crypt-like structures, with two type C aggregates between the crypt cross-sections. Each ROI contains similar numbers of each cell type and, hence, cell counts cannot distinguish the spatial structures. The Wasserstein distance measures overall similarity in cell distributions (e.g., similar distributions for A and B in all examples and differences for A and C in Exclusion). The QCM identifies positive and negative associations between cell pairs in small quadrats but fails to distinguish Exclusion from Architecture. The cross-pair correlation function (cross-PCF) quantifies correlations at a range of length scales. It identifies patterns where cell type A is uncorrelated with, clustered around, or excluded from C (compare *g*_*AC*_ (*r*) for Unstructured, Aggregation and Exclusion) and *g*_*AB*_(*r*) captures the periodically repeating clusters involving cell types A and B for Architecture. However, *g*_*AC*_ (*r*) does not distinguish between Exclusion and Architecture. The TCM identifies regions in which pairs of cell types are clustered or excluded, with persistent homology (PH) summarising these patterns. Although Exclusion and Architecture have similar *H*_0_ features (connected components), the *H*_1_ features (loops) in Architecture are more persistent, corresponding to crypt-like “islands.”

1. “Unstructured”, showing spatial independence between points with different labels;
2. “Aggregation”, reflecting dense clustering of some points into subregions of spatial co-location;
3. “Exclusion”, representing segregation of points with different labels into distinct subregions;
4. “Architecture”, representing higher order architectural structures such as colorectal crypts and/or immune aggregates. The stereotypical nature of these structures changes as neoplasia progresses.

While these examples are not exhaustive, they reflect an implicit hierarchy based on the spatial structure of the underlying tissue. We used them to assess the performance of our spatial descriptors in the identification of associations between points (representing cell centres) within different spatial contexts. Simple, non-spatial metrics, such as cell counts, are insufficient to distinguish the more complex spatial structures in Figure 1, as each dataset contains approximately equal numbers of points (or, equivalently, cells) with each label. The Wasserstein distance (*Wass*) (Villani 2009) describes the similarity between two cell distributions across a region of interest (ROI). It identifies exclusion between cell types (see high values of *Wass*_*AB*_ and *Wass*_*AC*_ in the “Exclusion” case), but fails to distinguish “Unstructured” and “Architecture” cases (both have low values of *Wass*_*AB*_, although their spatial structures differ). The quadrat correlation matrix (QCM) (Morueta-Holme et al. 2016) identifies cell count correlations within sub-regions, here 10% of the ROI width. At this length scale, the QCM identifies correlated and anti-correlated cell pairs in the “Aggregation”, “Exclusion” and “Architecture” cases. The cross-pair correlation function (PCF) (Baddeley, A., Rubak, E., and Turner, R. 2016) describes correlation between cell pairs across multiple length scales (determined by the parameter *r* in the output graphs). PCF values greater than one correspond to cell clustering, and values less than one to exclusion; the length scales associated with cell clustering and exclusion can be determined from the *r*-values of PCF maxima and minima. The “Unstructured” PCFs are close to 1 (consistent with complete spatial randomness, CSR), indicating point populations that are not correlated at any length scale (see also *g*_*AD*_ (*r*) in each case). The PCFs identify short-range aggregation between cell types A and B in “Aggregation”, “Exclusion” and “Architecture.” They also identify short-range exclusion between cell types A and C in “Exclusion” and “Architecture” but do not resolve differences in their organisational structures. The topographical correlation map (TCM) (Bull et al. 2024) identifies the presence and location of positive and negative co-localisation between cells in all examples. We use persistent homology (PH) (Robins 1999) to summarise peaks and troughs of the TCM, and represent their size, prominence and relationships to surrounding features as points in a persistence diagram (final column in Figure 1; see Methods). Importantly, while more complex spatial metrics may better distinguish spatial complexity, their interpretation becomes more difficult, suggesting that the statistic best suited to describe a given ROI represents a balance between interpretability and discriminatory ability.

Together, the data and analyses in Figure 1 illustrate the ways in which individual spatial descriptors characterise point associations across different synthetic models of tissue spatial organisation, and suggest an approach in which multiple spatial descriptors are used to analyse complex biological images.

### Multiple spatial patterns may be present in the same region of interest

Epithelial, immune and stromal spatial organisation and cellular architecture changes in the colorectum as invasive cancer emerges from benign polyps. Recognition of these changes in morphological patterns underpins the pathological diagnosis of cancer. Here, we applied our suite of mathematical tools to assess whether the altered cell associations that determine these fundamental histological changes could be quantitatively assessed in human tissue sections. We identified 43 carcinoma-in-adenoma specimens from the Oxford Rectal Cancer Cohort. These full tumour resection specimens include invasive cancer “caught-in-the act” of emerging from a pre-existing benign polyp precursor, representing two stages of colorectal cancer development within a single section. We identified markers for epithelium (E-Cad), a range of adaptive and innate immune populations (CD4, CD8, FoxP3, CD68, MPO) and mesenchymal cells (CD34, CD146, Periostin, Podoplanin, α-SMA), and undertook multiplex staining on two serial sections per sample. Cell markers were chosen following literature review to identify prognostically relevant cell types and enabled the analysis to focus on immune-stromal interactions (Zeng et al. 2017; Schimek et al. 2022; Rottmann et al. 2023; Galon et al. 2014; Ma et al. 2020; Wei et al. 2023; Lalos et al. 2021; Stzepourginski et al. 2017; Yamanashi et al. 2009; Kobayashi et al. 2022). The data were divided into 1000×1000 pixel (500μm x 500μm) ROIs, aligned using epithelial E-cadherin as a fiducial mark via a two-stage process as described in the methods. After additional quality control steps, this led to the generation of 10,027 well aligned ROIs across the cohort (see limitations of the work section). All mathematical methods were applied to each ROI within the dataset, to generate a set of spatial summary statistics (see Methods for details).

Our synthetic data (Figure 1) were designed to model tissue architecture relevant to the intestine, however, human tissues are complex, heterogenous and can exhibit multiple spatial organisational structures within an individual ROI. In adenoma, retention of crypt-based epithelial architecture typically resulted in restriction of immune and stromal cell interaction to interdigitating intercryptal ribbons (Figure 2A). In cancer, the loss of defined crypt structures is associated with formation of variable sized epithelial glands and the expansion of the desmoplastic stroma. These changes introduce spatial organisational variation across neoplastic phenotypes that could confound comparisons using single spatial descriptors. This is exemplified by looking at neutrophil-T helper cell interactions in representative ROIs from the same tissue sample (Figure 2A). In the adenoma region, the two cell types partly co-localise in the intercrypt mesenchyme, reflected by low Wasserstein distance (*Wass*_*NTh*_ = 84.75), short range clustering identified by the cross-PCF, the TCM and corresponding persistence diagrams (Figure 2B). In contrast, in this region of the carcinoma (Figure 2C), infiltration of T cells into cancer epithelial nests establishes neutrophil-T Helper cell exclusion, reflected by an increased Wasserstein distance (*Wass*_*NTh*_ = 136.45), weak exclusion in the cross-PCF at length scales up to 150 pixels, and the presence of ‘valleys’ in the TCM corresponding to persistent *H*_0_ features in the persistence diagram.

**Figure 2:**
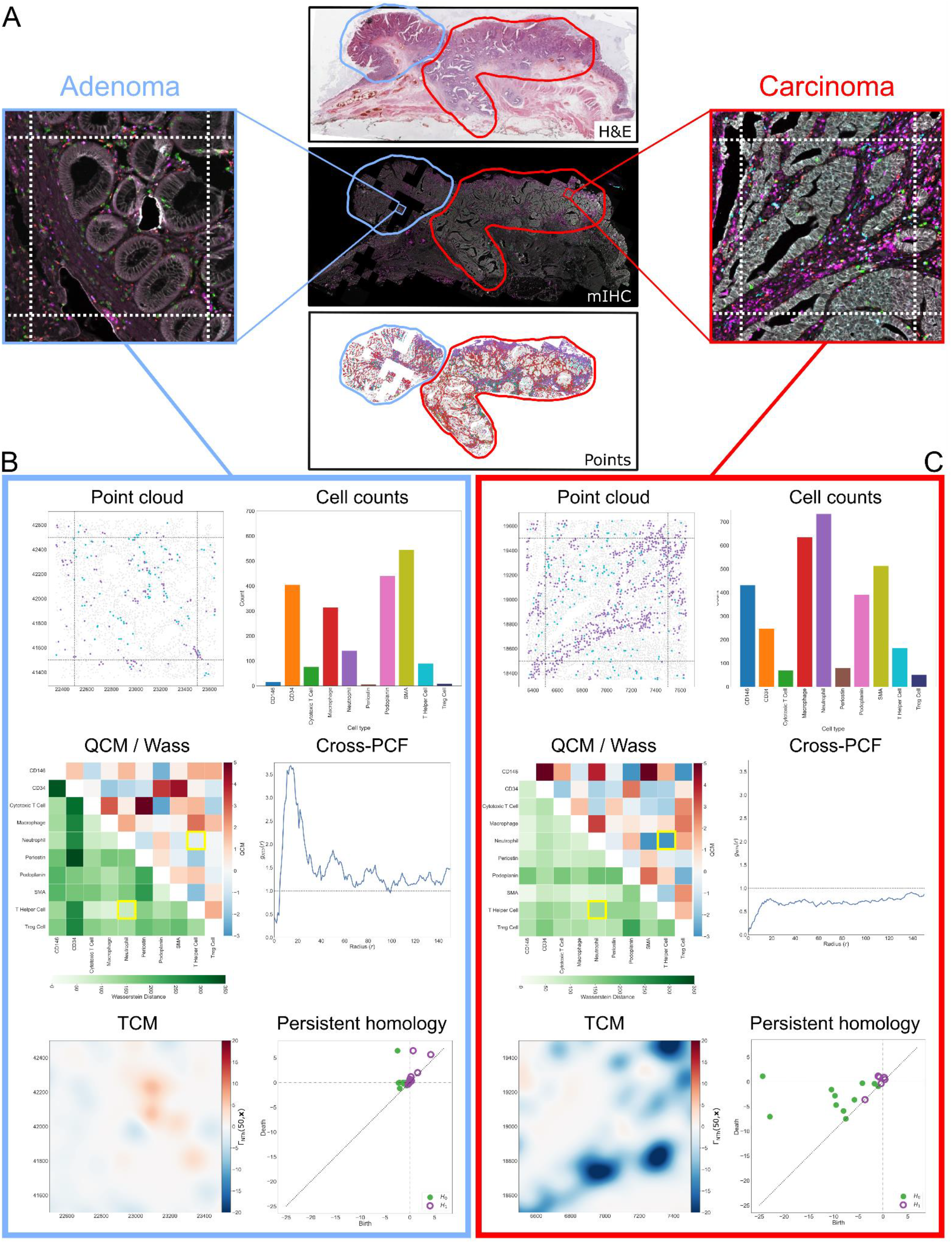
Multiple tissue organisational structures can be found within single regions of interest. Panel A: Representative patient sample, with H&E, multiplex IHC showing the immune panel (blue: regulatory T cell, red: macrophage, green: cytotoxic T cell, cyan: T helper cell, magenta: neutrophil, white: epithelium), and corresponding whole slide point cloud. Two representative ROIs annotated as adenoma (blue outline) and carcinoma (red outline) are highlighted. Panels B, C: Statistical descriptions of the ROIs in panel A. <u>Point cloud:</u> highlighting locations of neutrophils (purple) and T helper cells (cyan). In the adenoma these cell types are co-located in the stroma, surrounding crypt-like structures; in the carcinoma ROI, neutrophils are confined to the stroma whereas T helper cells also infiltrate the cluster of cancer cells. <u>Cell counts:</u> unprocessed cell counts within each ROI. <u>QCM / Wass:</u> Pairwise measurements of the Wasserstein distance (lower triangle) and QCM (upper triangle) between different cell populations, with neutrophils and T helper cell interactions highlighted. *Wass*_*NTh*_ is small in the adenoma, indicating similar cell distributions, and larger in the carcinoma, suggesting differing spatial distributions. <u>Cross-PCF:</u> Cross-PCFs between neutrophils and T helper cells, *g*_*NTh*_(*r*). In the adenoma ROI, strong correlation between the two cell types is observed at the length scale of approximately one cell diameter (10-30 pixels). In the carcinoma ROI, the two cell types are weakly anti-correlated at all length scales. <u>TCM:</u> TCMs between neutrophils and T helper cells, *Γ*_*NTh*_(50, ***x***). The carcinoma TCM shows several clusters of neutrophils, with no surrounding T helper cells (highly negative areas, shown in blue), while the adenoma TCM highlights several areas with weak positive and negative correlation. <u>Persistent homology:</u> Persistence diagrams showing *H*_0_ (filled circles, green; connected components) and *H*_1_ (open circles, purple; loops) features detected in the TCMs. Several carcinoma *H*_0_ features have large negative values for feature birth, highlighting the presence of multiple tissue regions in which neutrophils have strong local exclusion from T helper cells (negative values of the TCM). The adenoma *H*_0_ features generally have small birth and death values, with a small number of *H*_1_ features, suggesting a relatively flat TCM indicative of similar neutrophil and T helper cell distributions.

Together, these data demonstrate that the triangulation of multiple spatial descriptors across aligned whole tissue sections can account for heterogeneity in spatial organisational structures within an ROI and enable quantification of neoplastic phenotype specific differences in cell interactions.

### Combining complementary statistical features improves disease classification in individual patients

Our multiplex panel contained markers for 10 stromal and immune cell subtypes, yielding 100 possible ordered pairwise cell-cell combinations. We applied our suite of spatial descriptors and collated 14 features for each cell pair combination: Wasserstein distance (*Wass*_*ij*_), QCM (*QCM*_*ij*_), four scalars associated with the high and low values of the cross-PCF (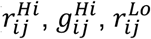 and 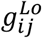) and eight that characterise the persistence diagrams in *H*_0_ and *H*_1_ (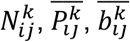 and 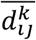 where *k* = 0,1).

Combination with cell counts produced 1410 features in each of the 10,027 ROIs across 43 carcinoma-in-adenoma samples. Taken together, these features define a profile that characterises spatial cell-cell relationships for each ROI. In this feature space, there is substantial overlap between adenoma and carcinoma across the cohort (see UMAP projection coloured by disease stage in Fig 3A; silhouette score = 0.015). However, at the individual patient level, we observed clustering of adenoma and carcinoma ROIs on a per-patient basis (see UMAP projection coloured by patient ID in Figure 3B, C), indicating heterogeneity in disease progression across the cohort.

**Figure 3:**
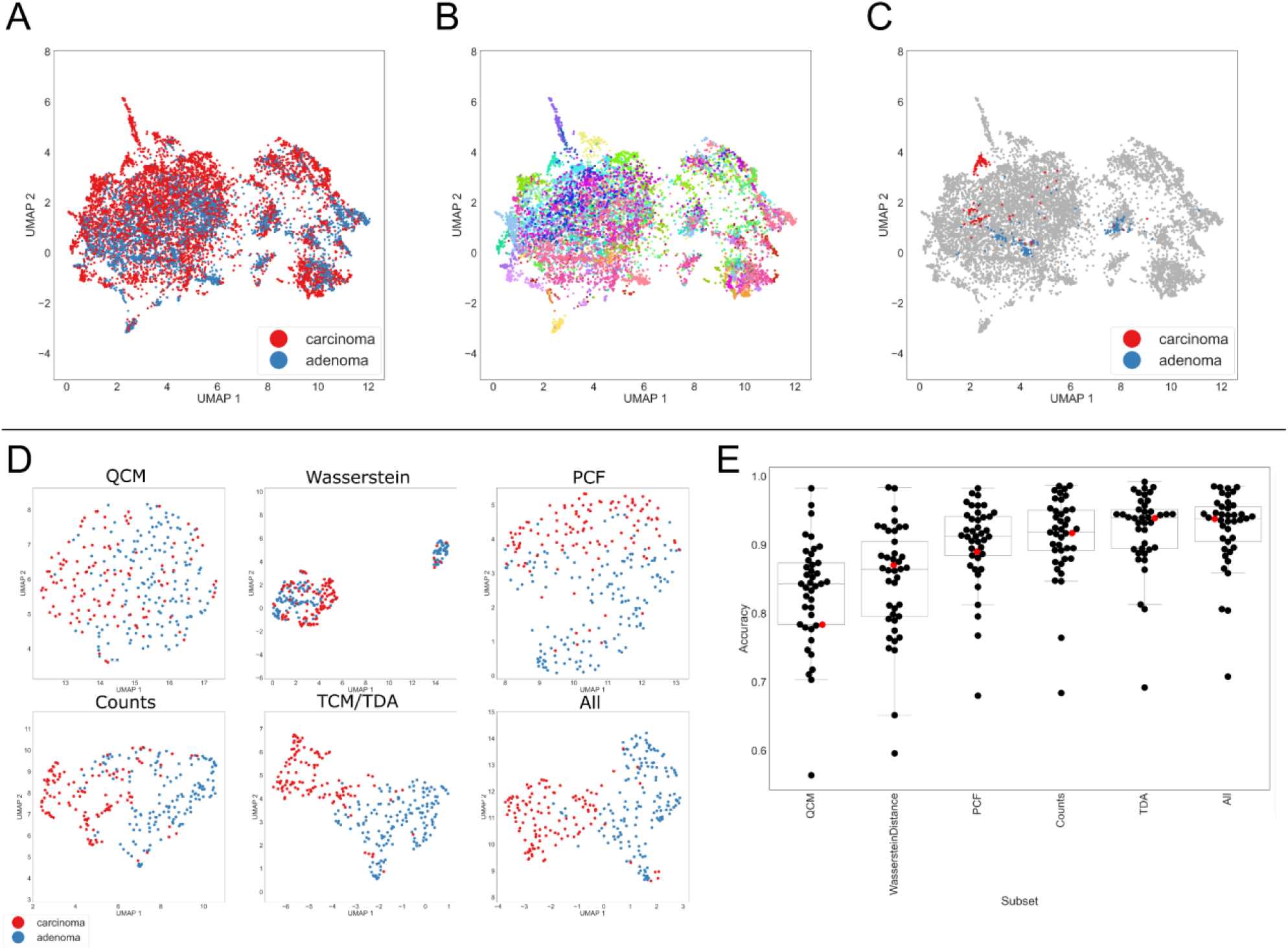
Inter-patient heterogeneity and the identification of stage-discriminatory cell interactions. Panels A-C: UMAP of all points in the 1,410 dimensional space defined by all summary features, coloured according to disease status (A), patient ID (B), and highlighting ROIs from a single patient (C). Panel D: series of UMAPs from a single patient, showing how the distinction between ROIs annotated as adenoma and carcinoma changes as we vary the statistics used to characterise the spatial distribution of the different cell types. Panel E: Using only the features specified, for each patient, 10 random forest classifiers were trained to distinguish adenoma from carcinoma on different 2:1 train:test splits of the data. Points represent the mean accuracy for each patient’s classifiers. Each feature generates reasonable classification, but using all features leads to the greatest classification accuracy. Red points represent the example patient in panels C and D. (Boxplots show quartiles and median, whiskers show samples within 1.5*IQR)

Motivated by this observation, we investigated the stage-discriminatory power of individual spatial descriptors on a per-patient basis. The degree of overlap in the spatial interaction profiles varied between methods. For a representative patient, visual inspection suggests that the QCM and Wasserstein metrics had lower discriminatory power than PCFs, cell counts, TCM/TDA and ‘All’ (see Figure 3D). This suggests that each subset of features provides complementary information about changes in cell-cell relationships between adenoma and carcinoma, and is consistent with the analyses in Figure 1 and Figure 2.

For each patient, we trained a series of Random Forest (RF) classifiers on subsets of the spatial descriptors in order to quantify their ability to characterise disease stages (after QC, 2 patients contained only carcinoma ROIs and were, therefore, excluded from RF analysis). Ranking the classifiers by their accuracy at predicting unseen ROIs from the same patient (see Methods) revealed a consistent trend across the patient cohort, with lowest accuracy for QCM (a=0.829) and Wasserstein (a=0.848) and highest accuracy for TDA (a=0.923) and ‘All’ (a=0.924) (Figure 3E).

Together, these data show that we can discriminate between adenoma and carcinoma ROIs within each patient with high accuracy, by using our suite of spatial descriptors which co-identify multiple neoplasia stage-discriminatory changes in specific cell-cell interactions.

### Multiple summary statistics co-identify the same patient-specific stage-discriminatory cell interactions

Next, we sought to identify the cell interactions that contribute most to stage discrimination in each patient. On a per-patient basis, we used a RF classifier trained on the full set of 1,410 features (‘All’ in Figure 3 and see Table 2 in methods). We use mean decrease in impurity (MDI, a measure of how well each feature improves classification; see Methods) to identify features with high importance in the classification. We report the 15 spatial features with highest MDI for two representative patients, patients A and B (Figure 4A and B), with bars coloured according to the statistical class from which the features are drawn (Figure 4C,D). For patient A, cell counts and TDA features relating to connected-components are prominent (Figure 4C) whereas in patient B, PCF and TDA features relating to loops are dominant (Figure 4D).

**Table 1:**
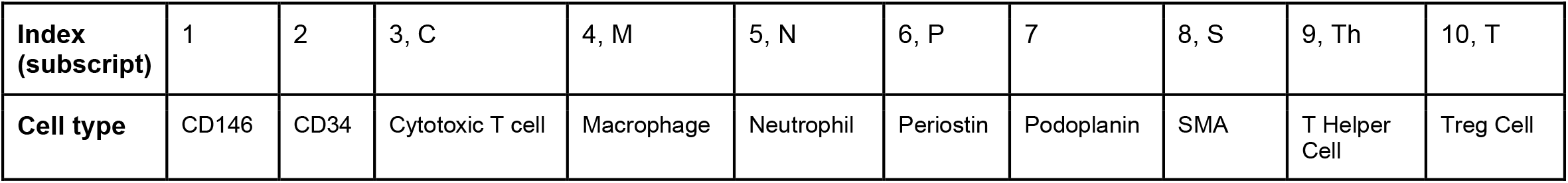
Indexing of cell types. Letters given are also used as subscripts in main text for notational convenience.

| Index (subscript) | 1 | 2 | 3, C | 4, M | 5, N | 6, P | 7 | 8, S | 9, Th | 10, T |
| --- | --- | --- | --- | --- | --- | --- | --- | --- | --- | --- |
| Cell type | CD146 | CD34 | Cytotoxic T cell | Macrophage | Neutrophil | Periostin | Podoplanin | SMA | T Helper Cell | Treg Cell |

**Table 2:**
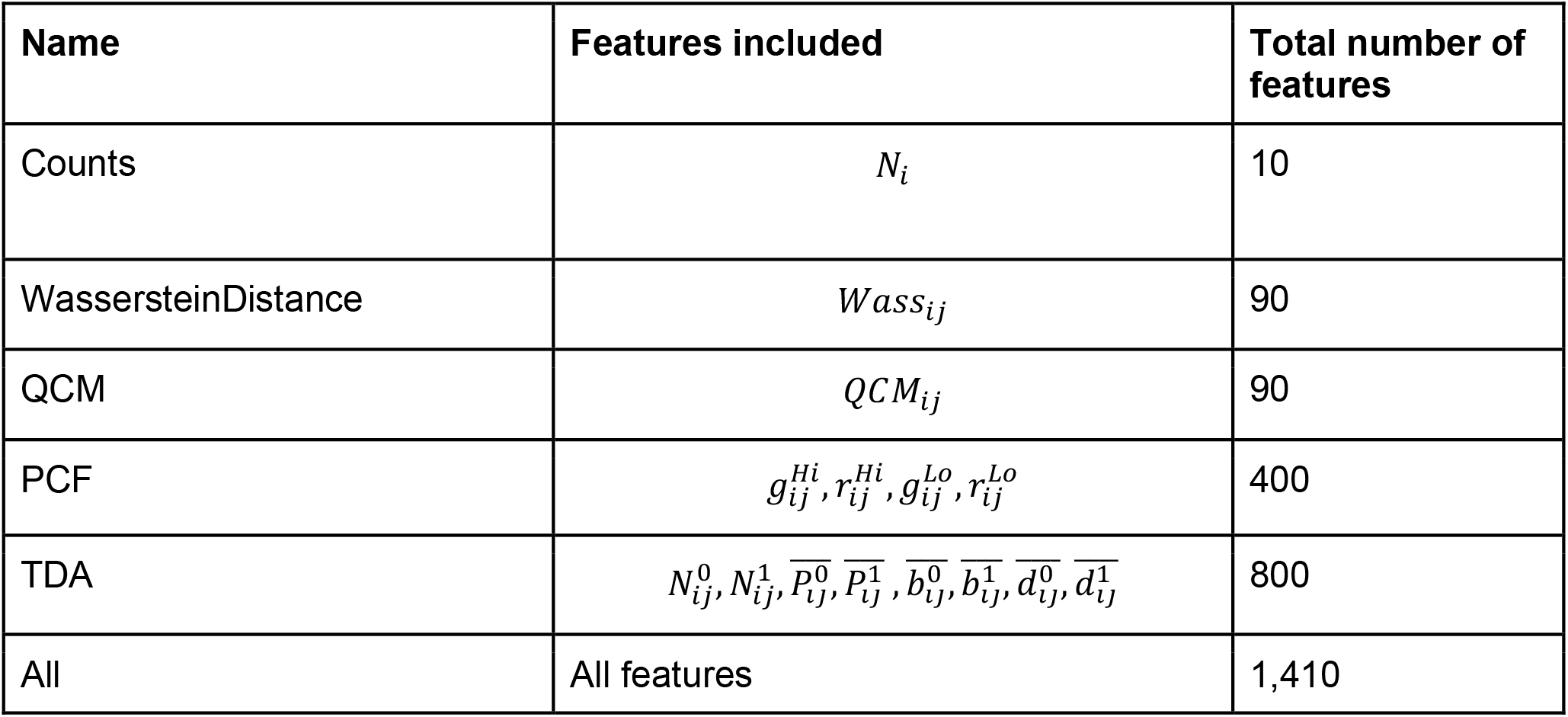
Subsets of features used to describe each ROI.

**Figure 4.**
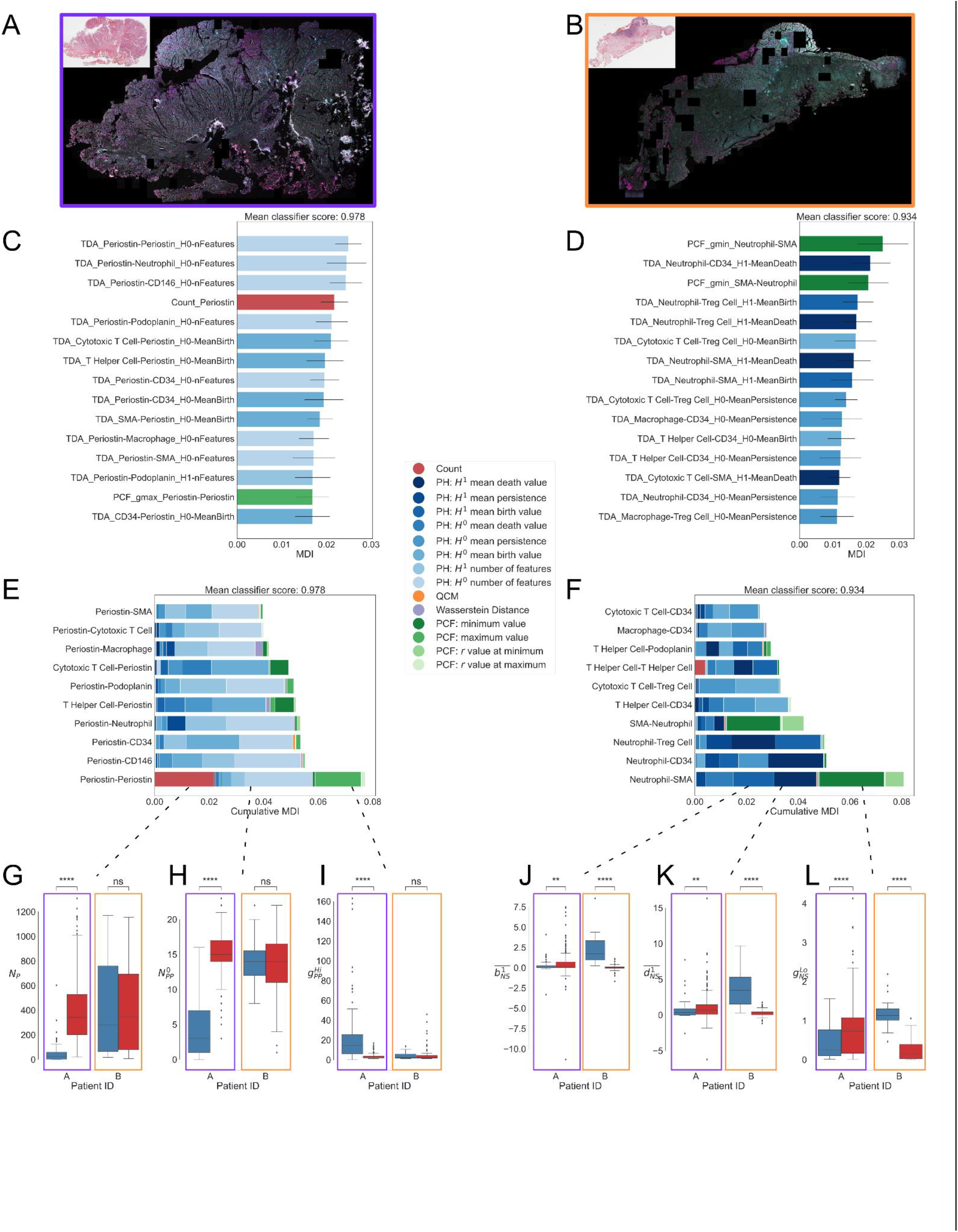
Multiple summary statistics co-identify the same stage-discriminatory cell interactions on a per patient basis. A, B: Parallel examples from two different patients (multiplex immune panel; blue: regulatory T cell, red: macrophage, green: cytotoxic T cell, cyan: T helper cell, magenta: neutrophil, white: epithelium; inset H&E). C, D: Top 15 most important features for discriminating adenoma/carcinoma in patients A and B, as determined by mean decrease in impurity in the corresponding random forest classifiers. E, F: Most important cell-cell pairs, according to cumulative MDI. G-L: Comparison of the extent to which 6 selected features are able to effectively discriminate between adenoma and carcinoma ROIs. Boxplots show median and quartiles, whiskers show 1.5*IQR. G-I: key features describing Periostin-Periostin interactions are discriminatory for patient A, but not significant for patient B; K-L: key features describing Neutrophil-SMA interactions are discriminatory for patient B, and less discriminatory for patient A (**: p<0.01, ****: p<0.0001, Mann-Whitney test, see Methods).

For each patient, we rank the MDI contribution of cell pairs to the classifier by aggregating spatial features associated with those cell-cell interactions (see Methods). Relationships involving periostin are prominent in patient A and not patient B. Conversely, neutrophils are prominent in patient B but not in patient A. The 10 most contributory cell pairs to the classifier are presented in the stacked plots in (Figure 4E and F). On closer inspection, three periostin-periostin features (the cell count, *N*_*P*_; the maximum cross-PCF value, 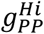 ; and the number of *H*_0_ PH features, 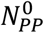) show significant variation between adenoma and carcinoma for patient A, but not for patient B (Figure 4G-I). In contrast, three statistics describing neutrophil-SMA interactions (the average birth and death values of *H*_1_ features,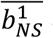 and 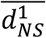; and the minimum value of the cross-PCF 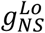) are strongly significant in patient B but less so in patient A (Figure 4J-L). Taken together, these results reinforce the hypothesis that the cell pairs that discriminate between adenoma and carcinoma are patient-specific.

### Patients can exhibit similar patterns of stage-discriminatory cell-cell interactions

Having established that triangulation of all developed spatial descriptors is effective at identifying stage-discriminatory cell interactions, we quantified the spatial relationships which characterise adenoma-to-carcinoma progression in each patient. We then compared the similarity of cell-cell interaction changes between samples by identifying patients with similar feature importance profiles. To each patient, we associated a 1,410 dimensional vector whose entries correspond to the MDI for each feature. We used principal component analysis (PCA) to identify combinations of features that most strongly define inter-patient variation (Figure 5A, B). We also identify groups of patients with similar stage-discriminatory cell relationship profiles. In particular, we note that patients with high PC1 are strongly influenced by interactions between periostin and a range of other cell types, while those with low PC1 are dominated by interactions involving innate immune cells. PC2 is influenced by periostin interactions, and interactions between SMA and immune cell populations (Figure 5B).

**Figure 5.**
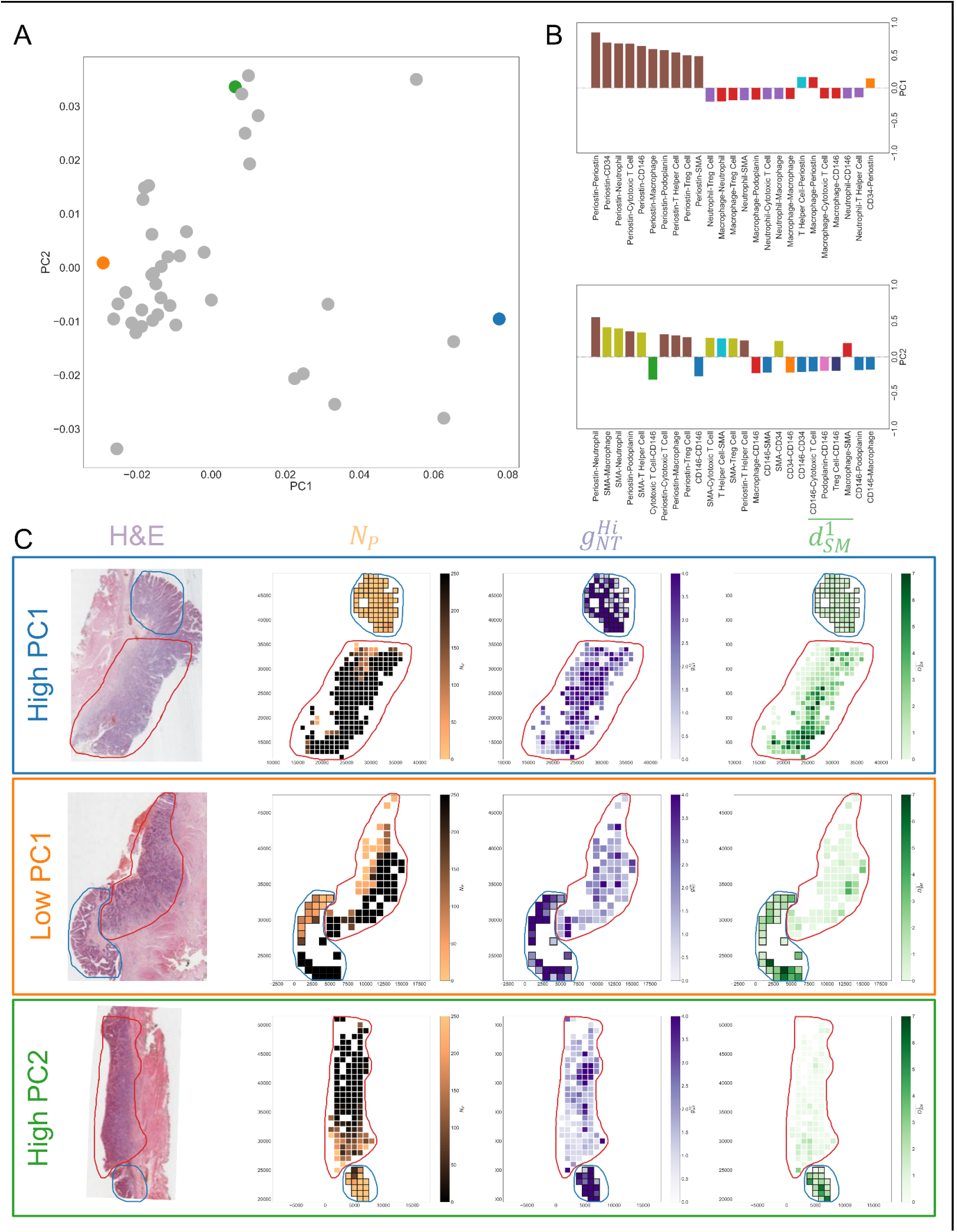
Stage-discriminatory interactions vary across the patient cohort. A: PCA shows grouping of patients with similar feature importances (variance explained: PC1 - 19.7%, PC2 - 8.0%). Coloured points correspond to the examples in panel C below. B: Interpretation of PC1 and PC2. Positive and increasing PC1 corresponds to increasing importance of Periostin-X interactions, while negative PC1 indicates Neutrophil-X or Macrophage-X. PC2 is linked with SMA and CD146 interactions. Bar colours correspond to the first cell type in each cell pair. C: Selected patients with positive/negative PC1 and high PC2 (see highlighted points in A), with ROIs coloured by the number of Periostin cells (*N*_*P*_), maximum cross-PCF between Neutrophils and T helper cells 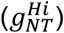, and the average death value of *H*_1_ features in the SMA-Macrophage TCM 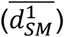. Adenoma (blue) and carcinoma (red) annotations are highlighted on all images.

To better understand these key cell interactions, selected features were mapped back onto the tissue for cross referencing with the tissue morphology from H&E images. We highlighted values of three different statistical features that represent important cell-cell interactions, and mapped them onto the tissue (Figure 5C). These three patient samples represent extreme values in the PCA (highlighted samples in Figure 5A). Notably, in samples with high PC1 (Figure 5C), the number of periostin positive cells (*N*_*P*_) is substantially different in the adenoma and carcinoma areas, while for samples from other regions of PCA space (Figure 5C, ‘Low PC1’ and ‘High PC2’) both the adenoma and carcinoma contain a mixture of regions with high and low periostin counts. More discriminatory for the low PC1 sample is 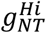 (the maximum value of the neutrophil-Treg cell cross-PCF), with higher values appearing in the adenoma than carcinoma. Finally, features involving SMA-macrophage interactions (here 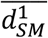, the average ‘death’ value of *H*_1_ features in the SMA-macrophage TCM) are more discriminatory for samples in the high PC2 region than for the other samples.

## Discussion

Advances in multiplex imaging have substantially progressed our understanding of cellular ecosystems, permitting detailed mapping of cell locations within a tissue and enabling the use of powerful tools to interrogate spatial relationships between cells of interest. These advances have the potential to transform the way we understand tissue architecture and disease pathology, but methodological development for quantitative analysis and interpretation of spatial data requires further research.

Numerous statistical tools have been used to analyse spatial biology datasets, but existing studies typically focus on a single spatial descriptor that is unable to resolve cellular relationships across the full range of length scales and tissue organisational structures seen in mammalian tissues. In this paper, we deployed a suite of applied statistical tools adapted from ecology, spatial statistics, measure theory, and topology, to determine whether different tools could identify the same biological cues within spatial data. Taken together, our results show how multiple tools can be used cumulatively to identify and cross-corroborate biological findings, leading to spatial analysis pipelines that are robust to the limitations of any particular method.

We focussed on statistical and mathematical metrics for quantifying spatial data, rather than leveraging artificial intelligence (AI) (Colling et al. 2019). Our methods are fully interpretable, and can be mapped back to specific properties of the underlying cell locations (such as those described in Figure 1). By contrast, existing AI approaches have great predictive power but require large training datasets and may not be applicable to data generated from different imaging platforms. Attempts to generate interpretable AI models are ongoing, however, identifying the spatial features they use to distinguish classes is challenging. The mathematical methods we use are better suited to exploratory analysis of cell-cell relationships, providing hypothesis-generating insights into how these relationships change with disease progression. In the longer term, convergence and cross-validation between the approaches has the potential to increase the power of both.

Our methods have broad applicability: the statistical tools can be used on cell coordinate maps, regardless of the spatial-biology platform used to generate them and the disease state under investigation. Here, we have used this approach to identify ways in which intercellular spatial relationships change during progression from benign adenoma to carcinoma in colorectal lesions. Our analysis shows that this tissue level change varies across a patient cohort, with inter-patient cellular ecosystem heterogeneity revealing variable cell interaction dynamics, despite cohort similarities in tumour stage (Stage 1), colonic position (rectum) and consensus molecular subtype (predominantly CMS4). By mapping the data in a high dimensional feature space, we can identify shared patterns of stage-discriminatory cell interactions in patient subgroups, indicating potential disease-defining biology that merits further investigation. Even with a comparatively small and economical marker panel, the power of spatial biological interrogation combined with combinatorial mathematical analysis can uncover variable cell-interaction pathways that underpin disease progression. Such insight may prove crucial in identifying the key, potentially druggable, signalling that mediates these interactions and in understanding variation in response to treatments such as immunotherapies. Together this work suggests that multiplex imaging and quantitative computational tools can be combined to empower pathologists to augment histological classification of pathological states, by identifying, assessing and contrasting dynamic cell interactions across disease contexts.

### Limitations of the study

This study has limitations, predominantly arising from the choice of spatial biology platform. Current spatial biology platforms are variably limited by ROI size, number of markers assessed, throughput capacity and expense. Given the number of patients and size of the whole tissue resections in this study, we needed multiplex capability with an unlimited staining aperture capable of generating whole slide images, and selected the Vectra Polaris platform which has a limited number of cell markers per run. To assess immune and stromal cell compartments, two consecutive whole slide images had to be registered to obtain cell coordinates, and this was achieved with a developed rigid alignment protocol, using epithelial cell staining on both panels as a fiducial mark (see Methods). However, unavoidable tissue and staining imperfections between sections meant that some quadrat regions failed our rigorous alignment quality control and were therefore not included in the analysis. Limited marker capacity also means that cell selection was based upon published evidence of cell type relevance in colorectal cancer. These limitations will be partly addressed as spatial platforms advance, with improving technologies (such as Xenium (10X Genomics) and CosMx (Nanostring)) allowing assessment of the spatially resolved expression of hundreds of genes. This will permit an unsupervised approach to cell interaction interrogation, free from marker selection bias. It may also provide some mechanistic insight into the cell signalling pathways that regulate key identified cell associations.

## Methods

### Experimental Model and Study Participant Details

Tissue sections were collected from Biobank project REC 11/YH/0020 (Oxford). Experimental slides were scanned and all subsequent work conducted on digital images which allowed slides to be rendered acellular in line with the Human tissue act requirements.

### Method Details

#### Multiplex imaging protocol

Akoya Biosciences OPAL Protocol (Marlborough, MA) was employed for multiplex immunofluorescence staining on FFPE tissue sections of 4-μm thickness. The staining was performed on the Leica BOND RXm auto-stainer (Leica Microsystems, Germany). A total of six consecutive staining cycles were conducted using primary antibody-Opal fluorophore pairs.

Multiplex IHC staining of epithelial, immune and, stromal cell–based biomarkers was performed to characterise the tumour microenvironment through disease evolution. The following markers were chosen to capture the high-level complexity of the intestinal tissue:

**Immune panel:** (1) <u>MPO</u> (Neutrophils, 1:4000, ab208670; Abcam)–Opal 540; (2) CD4 (T Helper Cells, 1:200, NCL-L-CD4-368, Leica;)–Opal 520; (3) <u>CD8</u> (Cytotoxic T Cells, 1:400, M7103; Dako)–Opal 570; (4) <u>CD68</u> (Macrophages, 1:400, M0876; Dako)–Opal 620; (5) <u>FoxP3</u> (Regulatory T Cells, 1:200, ab20034; Abcam)–Opal 650; and (6) <u>E-cadherin</u> (Epithelial Cells, 1:500, 3195; Cell Signalling)–Opal 690.

**Stromal panel:** (1) <u>PDPN (Podoplanin)</u> (Lymphatic Endothelium, 1:200, GTX21231; GeneTex)– Opal 540; (2) <u>CD34</u> (Mesenchymal Cell, 1:3000, ab81289; Abcam)–Opal 520; (3) <u>CD146</u> (Endothelial Cell,1:2000, ab75769; Abcam)–Opal 570; (4) <u>α-SMA</u> (Myofibroblasts, 1:1000, ab5694; Abcam)–Opal 620; (5) <u>Periostin</u> (Cell Adhesion Protein, 1:1000, ab227049; Abcam)– Opal 690; and (6) <u>E-cadherin</u> (Epithelial Cells, 1:500, 3195; Cell Signalling)–Opal 650.

The tissue sections were incubated with primary antibody for an hour, and the BOND Polymer Refine Detection System (DS9800, Leica Biosystems, Buffalo Grove, IL) used to detect the antibodies. Epitope Retrieval Solution 1 or 2 was applied to retrieve the antigen for 20 min at 100°C, in accordance with the standard Leica protocol, and thereafter each primary antibody was applied. The tissue sections were subsequently treated with spectral DAPI (FP1490, Akoya Biosciences) for 10 minutes and mounted with VECTASHIELD Vibrance Antifade Mounting Medium (H-1700-10; Vector Laboratories) slides. The Vectra Polaris (Akoya Biosciences) was used to obtain whole-slide scans and multispectral images at 20x magnification (0.5μm per pixel). Batch analysis of the multi-spectral images (MSIs) from each case was performed using inForm 2.4.8 software, and the resultant batch-analysed MSIs were combined in HALO (Indica Labs) to create a spectrally unmixed reconstructed whole-tissue image. Cell segmentation and phenotypic density analysis was conducted thereafter across the tissue using HALO. Regions corresponding to adenoma and carcinoma were manually annotated on each slide, and labelled point clouds consisting of cell centroid (x,y)-coordinates and assigned categorical cell type labels were extracted for registration and subsequent spatial analysis.

#### Point set registration via DARE (Density Adaptive point-set REgistration)

We registered point clouds using the Density Adaptive point set REgistration (DARE) algorithm (Lawin et al. 2018). DARE is designed to align point sets in which 1) sample points are not expected to be from identical locations, and 2) point density may vary substantially between point sets and across each point set. These properties render DARE suitable for aligning consecutive tissue sections using fiducial markers (E-Cadherin), since cells are expected to be from the same tissue areas but the same cell does not need to be present on both slides.

Further, DARE allows point set registration while accounting for regions of missing data that can be introduced during the scanning process (e.g. due to tissue damage or scanner failure), and for heterogeneity in detected cell density caused by variation in stain intensity across a slide In brief, the DARE algorithm is one of a class of probabilistic methods for aligning point sets, which identify a rigid geometric transformation (translation and rotation) that optimally aligns point sets by viewing them as samples from underlying continuous probability distributions (using, for example, a Gaussian Mixture Model) which should be aligned. The problem of aligning these distributions is viewed as an expectation-maximisation problem in which the likelihood of the points in one point set being obtained from the other probability distribution can be maximised. DARE modifies this approach to use a latent probability distribution that is weighted based on the local density of point observations, ensuring that areas in which points are densely clustered are weighted similarly to regions with sparse observations. This is a particular strength for registering serial tissue sections based on cell location data, since it ensures that registration relies on the underlying geometry of the tissue rather than variations in cell density artificially introduced by artefacts in cell classification induced by variations in tissue staining or classification, or by missing sections of tissue. Full mathematical details of the DARE algorithm can be found in (Lawin et al. 2018).

Note that throughout this manuscript we assume that cell locations from two consecutive slides occupy the same 2D space, which is misaligned. In reality, slide sections are separated in the z-stack by 4μm. When measuring cell-cell distances to calculate spatial metrics (described below), we neglect this 4μm offset, noting that the associated distance errors are below the minimum length scales considered in these metrics. In particular, the maximum error is 4μm (when two cells are deemed to be concentric, but are, in practice, separated by 4μm in the z-direction) and the measurement error introduced by a z-offset between slides shrinks rapidly as the distance in the (x,y) plane between the two points increases. (For example, points from separate slides that appear to be separated in the (x,y)-plane by 10μm are in reality separated by 10.77μm, and points with an apparent separation of 100μm are in reality 100.08μm apart.)

#### Global alignment

We apply the DARE algorithm using a two-stage procedure; first to generate an approximate global alignment, and then subsequently to each region of interest to ensure a precise local alignment. DARE permits some parameters to be varied to adjust the sensitivity of the algorithm; for global alignment, we randomly select 100,000 epithelial cells from each panel, and align using the parameters K = 150, num_iters = 1000, gamma = 0.005, epsilon = 1e-6, and num_neighbors = 20. Where global alignment was unsatisfactory (for example, when a substantial portion of tissue was missing from one slide), manual corrections to the rotation and translation were also applied to ensure a reasonable match between the tissue sections before local alignment.

#### Local alignment

After global alignment of the two panels, one slide is divided into a regular grid of non-overlapping 1000×1000 pixel regions of interest (ROIs). For each of these regions of interest, a local correction to the global alignment is applied, calculated from the 5000x5000 pixel region surrounding the ROI centre (i.e., alignment is based on the ROI together with a surrounding 2000 pixel buffer). Local alignment is only attempted if there are a minimum of 50 E-Cadherin positive cells on each panel within a 2000 pixel square centred at the ROI centre, and a minimum of 5 E-Cadherin positive cells on each panel within the ROI, otherwise the ROI is discarded from further analysis. Details of further quality control of the local alignment using spatial metrics is provided below in the section “*Region of Interest (ROI) acceptance criteria*”.

#### Overview of spatial metrics

We perform spatial analysis of point populations using four mathematical methods that derive from distinct subject areas: optimal transport (Wasserstein distance), ecology (quadrat correlation matrix, QCM), spatial statistics (cross-pair correlation function, cross-PCF), and topology (persistent homology). These methods quantify different features of spatially-resolved point clouds involving two cell types, with labels *C*_1_ and *C*_2_ say, and are briefly introduced in this section before being defined in detail below.

The Wasserstein distance describes the similarity of the spatial distributions of two sets of points, across an ROI, and is a powerful tool to measure spatial variations in local cell population densities within the domain. However, as a measure of similarity, it is unable to distinguish complex spatial interactions, such as spatial correlation between cell types separated by distances greater than *r* = 0, or higher order patterns such as loops or other topological features (see Figure 1, main text). The QCM resolves differences between two sets of points at length scales below the domain size, identifying correlation between cell counts of different types while accounting for the underlying spatial structure of the tissue by using random relabelling to generate a null distribution against which observations can be compared. However, the QCM relies on a specified length scale on which cell counts are observed, and is unable to quantify more complex spatial structures or patterns at length scales beyond the one chosen.

The cross-PCF quantifies correlation between pairs of cells that are separated by a distance r for a range of values of r, to identify length scales on which the two cell populations are positively or negatively correlated. The cross-PCF can identify the presence of clusters of groups of cells, or exclusion between cells of different types. However, the cross-PCF is unable to identify higher order anisotropic structures such as loops or patterns, particularly in the presence of noisy biological imaging data (Vipond et al. 2021). The most complex method considered here, persistent homology, is a form of topological data analysis (TDA) which is designed to identify topological structures such as “connected components” (groups of cells of the same type in close proximity) or “loops” (groups of cells which form connected components around a “void” from which they are excluded). While recent advances permit persistent homology to be applied directly to point patterns containing multiple cell types (di Montesano et al. 2024; Stolz et al. 2023), in this work we first pre-process the data to generate the topographical correlation map (TCM) (Bull et al. 2024), a spatially-resolved summary of local interaction between cells of types *C*_1_ and *C*_2_, and use persistent homology to summarise the higher order structures (loops and connected components) in this data.

#### Wasserstein Distance

The p-Wasserstein distance (or Kantorovich–Rubinstein distance), which for convenience we refer to throughout the main text as the Wasserstein distance, is a standard method from the field of optimal transport defining a distance between two probability distributions (Villani 2009). Intuitively, the Wasserstein distance yields the minimum cost to transform one distribution onto another. Consider the discrete measures *μ* = (*μ*_1_, …, *μ*_*n*_) and *v* = (*v*_1_, …, *v*_*m*_) with spatial positions ***x*** = (***x***_1_, …, ***x***_*n*_) and ***y*** = (***y***_1_, …, ***y***_*n*_) within the domain Ω.

The p-Wasserstein distance can be defined as:

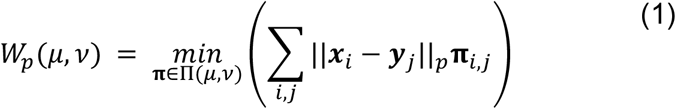

where ∥ ⋅ ∥_*p*_ is the *l*_*p*_-norm and

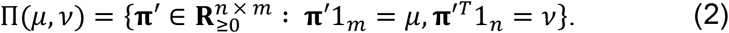

Namely, **π** provides the optimal transport plan to transform μ onto *v* with minimal work. The Wasserstein distance is a formal mathematical norm and therefore is positive (*W*_*p*_(*μ*, *v*) ≥ 0 and *W*_*p*_(*μ*, *v*) = 0 if and only if *μ* = *v*) and symmetric (*W*_*p*_(*μ*, *v*) = *W*_*p*_(*v*, *μ*)) (Santambrogio 2015). Throughout this study we use p=2 and therefore we denote *W*_2_(*μ*, *v*)= *W*(*μ*, *v*). Moreover, we assume that each sampled points are in 2D domains that are uniformly weighted, i.e., ***x***_*i*_, ***y***_*j*_ ∈ **R**^2^ where 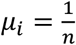 and 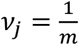 for all 1 ≤ *i* ≤ *n* and 1 ≤ *j* ≤ *m*.

Solving for **π** is computationally inefficient for distributions in more than one dimension and subsequently we employ the sliced modification of the Wasserstein distance (Bonneel et al. 2015). That is, the distributions *μ* and *v* are projected onto a 1D line that is drawn through the barycentre at an angle *θ*∼*U*[0,2*π*) and the 1D Wasserstein distance then computed,

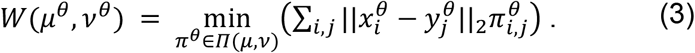

The sliced Wasserstein (*SW*) is then defined as the root average of the 1D Wasserstein distance of the sampled projections,

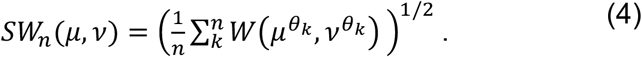

Critically, *lim*_*n*→∞_ *SW*_*n*_(*μ*, *v*) = *W*(*μ*, *v*) (Xi and Niles-Weed 2022). In this study, the sliced Wasserstein distance is calculated using the `ot.sliced_wasserstein_distance()` function from the Python Optimal Transport (POT) module (Flamary et al. 2021) with *n* = 50 (the module default), taking ***x*** and ***y*** as the spatial positions of the cell type pair *C*_*i*_ and *C*_*j*_, respectively.

#### Summary statistics

Since the sliced Wasserstein distance returns a scalar to describe the similarity of two point patterns, we include it directly in the combined feature vector for subsequent analysis (see below).

- *Wass*_*ij*_ = *SW*_50_(*μ*_*i*_, *v*_*j*_), the sliced Wasserstein distance for cell populations *i* and *j*, where *μ*_*i*_ and *v*_*j*_ denote the distributions of the two populations respectively.

#### Quadrat Correlation Matrix (QCM)

The Quadrat Correlation Matrix (QCM) describes correlations between the counts of different cell types within squares, or ‘quadrats’, of a specified edge length. First defined in an ecological context (Morueta-Holme et al. 2016), it has since been used to analyse correlations between cell types in digital pathology images of colorectal cancer multiplex immunohistochemistry slides (Gatenbee et al. 2022) and imaging mass cytometry images of Covid-19 lung sections (Weeratunga et al. 2023). The QCM identifies statistically significant co-occurrences within a ROI between cells of different types, comparing the strength of the observed correlation between a given pair of cell types against the expected correlation that would be observed if cell labels were assigned randomly. It relies on a choice of length scale which defines the size of each quadrat, which we take at 10% of the ROI size following previous work (Bull et al. 2024; Weeratunga et al. 2023). We therefore divide each 1000 x 1000 pixel ROI (500*μ*m x 500*μ*m) into a regular grid of 100 non-overlapping quadrats of size 100 x 100 pixels each (50*μ*m x 50*μ*m). The QCM is then constructed as follows:

1. A matrix ***O*** of observed cell counts is generated from the cell data. Each entry *O*_*ij*_contains the number of cells of type *i* found in quadrat *j*, for 1 ≤ *i* ≤ *n* and 1 ≤ *j* ≤ *m* where *n* is the number of different cell types and *m* is the number of quadrats.
2. A null distribution of 1000 (*n x m*) matrices ***N***^1^, …, ***N***^1000^ is generated by shuffling the cell labels of the points in the ROI. This is achieved by repeatedly resampling the entries of ***O*** while keeping the row and column sums constant, following the rules:

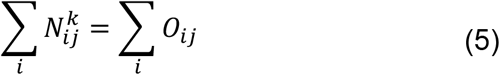

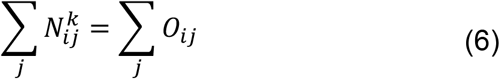 Computationally, we construct each matrix ***N***^*k*^ by repeatedly sampling two random rows (*a, b*) and columns (*c, d*), together with a random integer *q* sampled from the interval 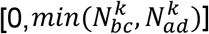, and then setting 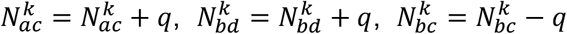 and 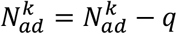. This process is repeated 10,000 times for each *k* to ensure that the matrix is well-shuffled.
3. Partial correlation matrices ***P***_***O***_, and 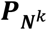 are generated for ***O*** and for each of the ***N***^*k*^.
4. The Standard Effect Size (SES) describing the strength of the observed correlation of cell types *i* and *j* relative to correlations from the null distribution is calculated using the mean, *μ*, and standard deviation, *σ*, of the null distribution, according to the formula:

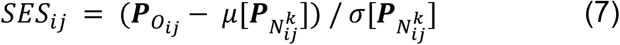
5. Significant SES can be identified by calculating a 2-tailed p-value for each pair of cell types and applying a Benjamini-Hochberg correction to account for multiple testing, with false discovery rate FDR = 0.05. The QCM is obtained by setting non-significant entries of SES to 0.

#### Summary statistics

For each ordered pair of cell types *C*_*i*_ and *C*_*j*_, we add the corresponding entry *SES*_*ij*_ defined in Equation (7) to the feature vector used for classification, noting that this means no significance testing is associated with the entries in the feature vector described as coming from the “QCM”. We also note that the QCM is not defined when *C*_*i*_ = *C*_*j*_.

- *QCM*_*ij*_ = *SES*_*ij*_

#### Cross-pair correlation function (cross-PCF)

The cross-pair correlation function (cross-PCF) is a spatial statistic which describes how frequently pairs of points with cell types *C*_1_ and *C*_2_ are observed to be separated by a distance *r* in a region of interest, relative to the number of such observations expected under a statistical null hypothesis of complete spatial randomness (CSR). Consider a set of cells in a region of interest (ROI) with area *A*, which we take to be a closed and bounded subset of ℝ^2^.

Mathematically, we consider each cell *i* to be represented by its centroid, a point ***x***_*i*_ = (*x*_*i*_, *y*_*i*_), together with a categorical mark *c*_*i*_ which describes its assigned cell type. For convenience, we introduce an indicator function, 𝕀, to determine whether the mark *c*_*i*_ matches some specified category *C*:

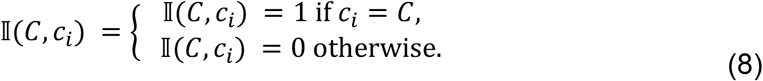

Similarly, it will be necessary to measure whether two points are separated by a distance “close” to *r*; in this work we consider a series of annuli whose inner radii are at discrete values *r*_*k*_ and of width *dr*, following our previous usage (Bull et al. 2024), although we note that the cross-PCF can be defined using a wide range of other distance kernels (Baddeley, A., Rubak, E., and Turner, R. 2016). We formalise this using an indicator function, *I*_[*a,b*)_(*r*), which equals 1 if *r* ∈ [*a, b*) and is 0 otherwise.

Mathematically, the cross-PCF for empirical data can be defined as:

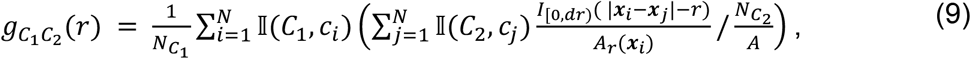

where 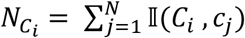 is the number of points in the ROI with cell type *C*_*i*_, *N* is the total number of cells in the domain (ROI), and *A*_*r*_(***x***) is the area of the intersection between the domain and an annulus centred at ***x*** with inner radius *r* and outer radius *r* + *dr*. This definition of *A*_*r*_(***x***) provides an implicit boundary correction for points near the edge of the ROI which permits the cross-PCF to be well-defined in a ROI with non-rectangular boundaries, but we note that many other alternative boundary correction terms could be used (Bull et al. 2024; Baddeley, A., Rubak, E., and Turner, R. 2016). Practically, we calculate 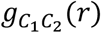 at a series of discrete values *r*_*k*_, such that *r*_0_ = 0 and *r*_*k*+1_ = *r*_*k*_ + *δr*. For all cross-PCF plots shown in this paper, we use the parameters *dr* = 10 and *δr* = 1, and consider values of *r*_*k*_ up to *r* = 150. To improve computational speed, when calculating cross-PCFs on all ROIs we use δ*r* = 5.

#### Summary statistics

For a given ordered pair of cell types, (*C*_*i*_, *C*_*j*_), we summarise the cross-PCF 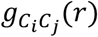 using four scalar summary statistics designed to capture the maxima and minima of the cross-PCF, detailed below:

- 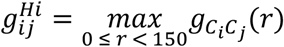, the highest observed value of 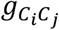.
- 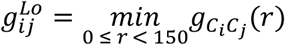, the lowest observed value of 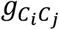.
- 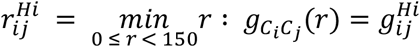, the *r* value that maximises 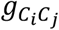.
- 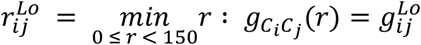, the *r* value that minimises 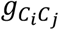.

### Topographical Correlation Map (TCM)

The Topographical Correlation Map is a variation on the general class of local indicators of spatial association (LISA) (Anselin 1995) that identifies the strength of local positive or negative correlation between pairs of cell populations. The TCM uses a linearised mark to represent contributions to positive or negative correlation of each cell in the cross-PCF, which permits marks to be summed to provide a spatially-resolved overview of which parts of a ROI contribute most strongly to the cross-PCF. To define the TCM between cells of types *C*_1_ and *C*_2_, *Γ*_12_(*r, x, y*), we first assign to each cell of type *C*_1_ a mark *m*_12_, which represents the cumulative sum of that cells contributions to the PCF up to radius *r* (or, alternatively, the value of Ripley’s K function (Baddeley, A., Rubak, E., and Turner, R. 2016; Ripley 1977) at *r*):

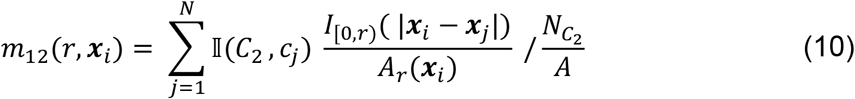

where the components of the equation are as defined in the previous section. In comparison to Equation (9), the interpretation of *m*_12_ is the same as for the cross-PCF, with *m*_12_ > 1 indicating clustering and *m*_12_ < 1 indicating exclusion.

Equation (10) defines *m*_12_ only at the location of cells of type *C*_1_. To facilitate the extension of Equation (10) to the full spatial domain, we first normalise this mark to ensure that multiple marks can be summed without distorting their interpretation:

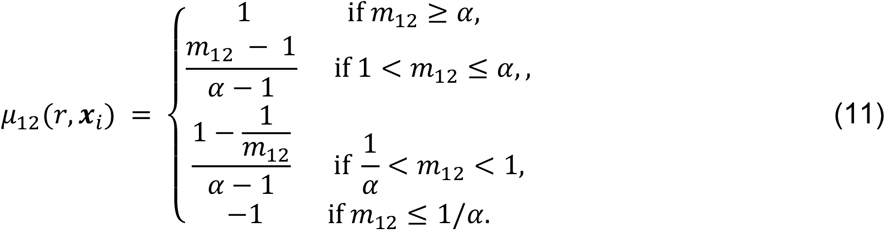

The purpose of this renormalisation is to ensure that *μ*_12_ is bounded between 1 and -1, with these representing the maximum resolution of positive and negative correlation between cells of types *C*_1_ and *C*_2_ respectively, determined by the parameter α. Within this range, positive and negative values of *μ*_12_ can be interpreted as an equal strength of correlation with opposite direction; i.e., *μ*_12_ = 0.5 indicates that cells are clustered with strength 0.5*α* while *μ*_12_ = −0.5 indicates exclusion with strength 0.5*α*. Further details and in depth discussion of this normalisation stage can be found in (Bull et al. 2024).

The normalised spatial marker, *μ*_12_, is defined only at locations where a cell of type *C*_1_ is present. In order to visualise the combined influence of all cells on local correlation across the domain, we centre a Gaussian kernel with standard deviation *σ* and scaled by *μ*_12_(***x***_*i*_) above each cell of type *C*_1_. The TCM, *Γ*_12_(*r*, ***x***), is then obtained by summing these kernels over all cells with mark *C*_1_:

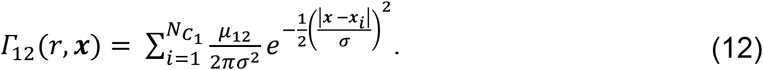

Note that the TCM is not symmetric - i.e., *Γ*_12_(*r*, ***x***) ≠ *Γ*_21_(*r*, ***x***), since the locations of the Gaussian kernels are based on the first cell population of interest. Following previous work, we take *σ* = *r μm* and *α* = 5, see (Bull et al. 2024) for further details of these choices and examples of the TCM applied to different patterns of cell distributions.

#### Summary statistics

The TCM, *Γ*, is a complex, spatially resolved, output, so we do not generate summary statistics for an ROI directly from it. Instead, we first apply topological data analysis (TDA) to summarise the TCM, which is described in the following section.

### Topological Data Analysis

To compare TCMs across different ROIs, we use topological data analysis (Ghrist 2008; Carlsson 2009; Edelsbrunner, Letscher, and Zomorodian 2000; Edelsbrunner and Harer 2006; Robins 1999), a field that studies shape and connectivity in data. Specifically, we applied persistent homology (PH), a robust and multiscale method in topological data analysis that allows tracking of topological features such as connected components, *H*_0_, and loops, *H*_1_, across a range of thresholding scales. As an input to PH we used matrices containing the TCM discretised evenly across 100 values spanning the *x* and *y* scales (i.e., a (100×100) matrix), describing a spatial correlation heatmap of cell types *C*_*i*_ and *C*_*j*_ in the ROI. We threshold each heatmap across *Γ* values starting from the lowest value in the heatmap to create a stack of images, a so-called sublevel set filtration (Edelsbrunner and Harer 2006), up to the highest value (see Figure 6). In each step of the filtration, we interpret the thresholded TCM as a 2D surface. Note that since the TCM can have negative entries, the thresholding values can be negative. PH allows us to observe the changes in the connected components and loops of this surface across the filtration.

**Figure 6.**
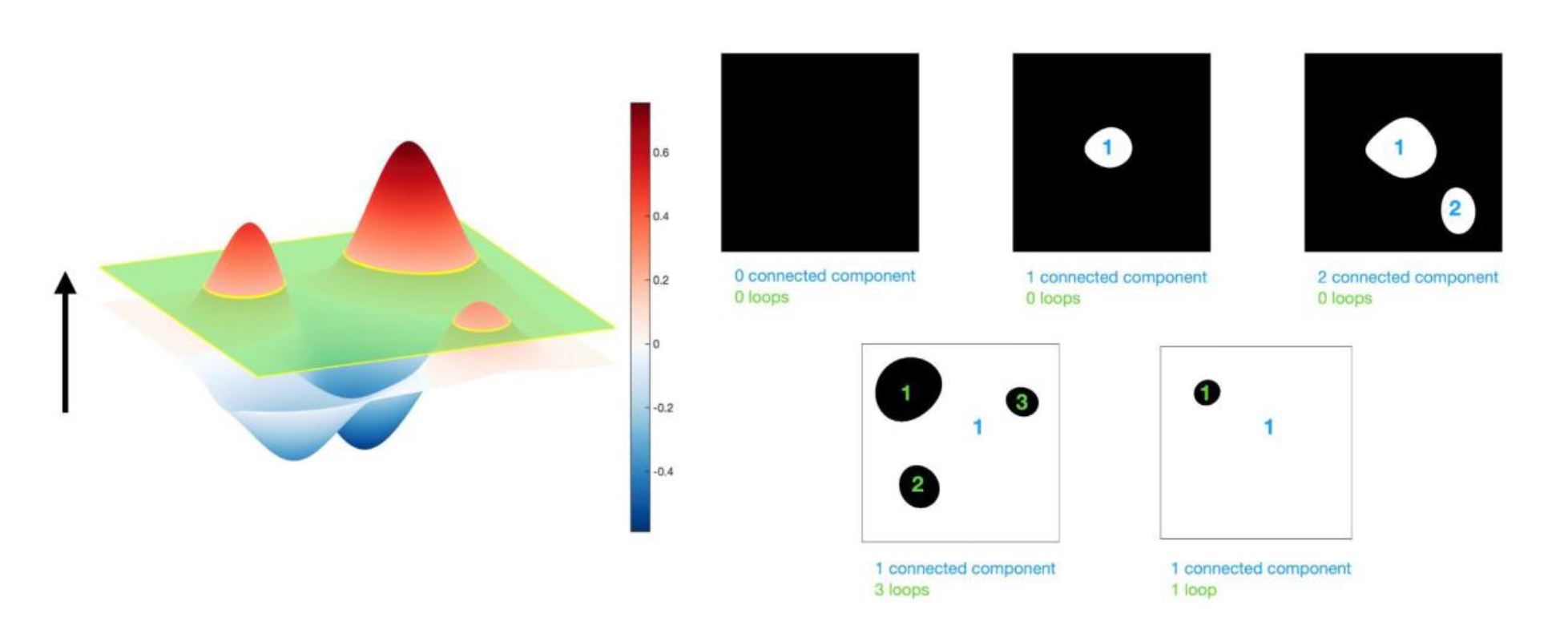
Schematic showing a sublevel set filtration applied to a TCM heatmap. Since the TCM contains both negative and positive values, the filtration parameter begins from the most negative value and moves up the TCM surface. As the filtration parameter increases, intersections with the “troughs” of the surface generate connected components in the mask of pixels included at that stage of the filtration (white). “Peaks” of the TCM appear as loops in the filtration mask. When the filtration parameter reaches the top of the TCM surface, all pixels are included in the filtration and the analysis stops.

A range of methods exist to visualise the output from PH; we use persistence diagrams, a method through which distinct PH features are plotted as points. In a persistence diagram, the x-coordinate corresponds to the values of the threshold parameter at which they are “born” (i.e. the value at which a connected component or loop first appears in the filtration) and the y-coordinate corresponds to the values at which they “die” (i.e., the value at which a connected component merge with other features or a loop is fully covered by pixels). We present *H*_0_ (connected components) and *H*_1_ (loops) features on the same diagram, using filled green circles to represent *H*_0_ features and unfilled purple rings to represent *H*_1_ features. The persistence of a feature is defined as the difference between the corresponding death and birth values, i.e. the range of threshold values across which the feature persists. For *H*_0_ features in the TCMs, i.e. connected components, birth values indicate troughs while their persistence is a measure of the depth of these troughs and how isolated they are. For *H*_1_ features, i.e. loops, death values point to peaks and their persistence indicates the height of the peak. In both cases, the filtration values of the TCM at the top or bottom of the peak (death in *H*_1_ or birth in *H*_0_) are related to the correlation strength of the feature (and whether it is positive or negative).

Let *X* denote our input heatmap, representing the function *X* : ℝ^2^ → ℝ, where each point in the heatmap has a corresponding co-localisation value between two distinct cell-types, *C*_*i*_ = *C*_*j*_.

The sublevel set filtration is constructed by sequentially considering sublevel sets *X*_*t*_ at consecutive thresholds *t* defined by *X*_*t*_ = {(*x, y*) ∈ *X* | *X*(*x, y*) ≤ *t*}. The filtration parametrized by increasing values of *t* gives rise to inclusion maps between subsets 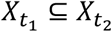 where *t*_1_ ≤ *t*_2_. We can interpret the sublevel set *X*_*t*_ at filtration value *t* as a simplicial complex, i.e. as a collection of nodes (0-simplices), edges (1-simplices), and triangles (2-simplices). We add each pixel as a node and add edges between the 8 pairwise adjacent nodes in the 2D Moore neighbourhood. We interpret three pairwise connected nodes as a triangle. We can now compute the homology groups *H*_0_(*X*_*t*_) and *H*_1_(*X*_*t*_) - representing connected components and loops at level *t* of the filtration - via boundary maps ∂_*k*_, which are linear maps sending *k*-dimensional simplices in a simplicial complex to their boundary, i.e. the sum of its (*k* − 1)-dimensional faces. Here, we only consider boundary maps ∂_2_, ∂_1_ and ∂_0_, i.e. *k* = 0,1,2. The boundary of a boundary is always empty, i.e. ∂_*k*_ ∘ ∂_*k*+1_ = 0. Computing the kernel *Ker*(·) and image *Im*(·) of the boundary maps we obtain the vector spaces *H*_0_(*X*_*t*_) and *H*_1_(*X*_*t*_) as follows:

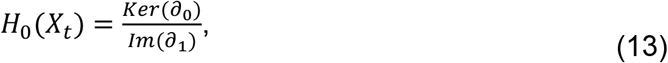

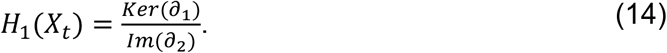

The inclusion maps between the sublevel sets 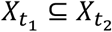 induce maps between their homology groups 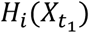 and 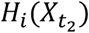 which allow the construction of a persistence diagram to track the elements of the vector spaces across the filtration. For a more detailed description of the topological background, see (Ghrist 2008; Carlsson 2009; Edelsbrunner, Letscher, and Zomorodian 2000; Edelsbrunner and Harer 2006; Hatcher 2015).

We compute the sublevel set filtration using the software Ripser (Bauer 2021) within a Python implementation. The resulting barcodes can be summarised by a range of interpretable metrics to allow statistical analysis. Note that there are many possible choices for such metrics, as well as various vectorisation methods (Ali et al. 2023; Adams et al. 2017) including persistence landscapes (Bubenik 2015).

#### Summary statistics

For a given ordered pair of cell types, (*C*_*i*_, *C*_*j*_), we summarise the persistence diagrams calculated on a TCM using 8 different summary statistics, briefly described below. These reflect the number of features in the diagrams 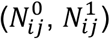, mean persistence for each feature 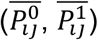, and the mean birth and death values 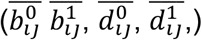. In this notation, the superscript indicates whether the value was obtained from an *H*_0_ or *H*_1_ diagram.

- 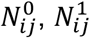= Number of features
- 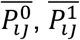= Mean persistence
- 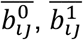 = Mean birth
- 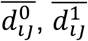 = Mean death

### Summary feature vector for statistical analysis

We use the summary statistics above to define a feature vector for each region of interest (ROI), which represents each ROI as a point in a high dimensional space. Across the two panels, we have a total of 10 cell types of interest (excluding epithelial cells):

Based on these indices, we define for each ordered cell pair (*i, j*):

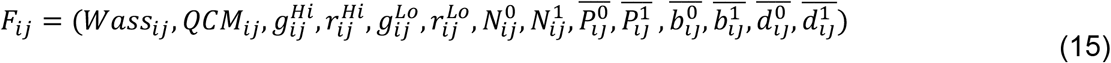

Combining all 100 possible ordered combinations of cell-cell interactions with the 10 cell counts, the final feature vector defines a 1,410 dimensional space forming the basis for subsequent analyses:

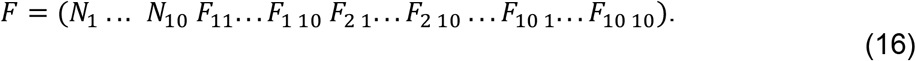

In the analysis for Figure 3, we consider spaces defined by subsets of this feature vector. The relevant entries in Equation (15) for each description in Figure 3 are shown in the following table, where *i* and *j* include all values from 1 to 10:

### Quantification and Statistical Analysis

#### Region of Interest (ROI) acceptance criteria

Each annotated adenoma / carcinoma image was completely tiled into non-overlapping 1000 x 1000 pixel ROIs (500*μ*m x 500*μ*m), based on the point cloud of extracted cell centres on the immune panel. Point clouds from the second panel were aligned with each ROI through the process described above (see “*Point set registration via DARE (Density Adaptive point-set REgistration)”*, Methods), based on a first approximate global registration followed by more refined local registration for each ROI.

A total of 20,212 ROIs contained at least one cell. For each of these, the alignment quality was automatically assessed using two complementary methods to compare the distribution of the fiducial epithelial marker on the two panels, based on the Wasserstein distance and cross-PCF. First, we calculated *g*_*Epi* 1,*Epi* 2_(*r*), the cross-PCF between the epithelial markers on each panel, to identify whether one panel was translated from the other. Such a shift would lead to strong correlation between the fiducial markers at a distance greater than *r* = 0, which would identify the amount by which the slides were misaligned. Any ROI for which the maximum value of *g*_*Epi* 1,*Epi* 2_(*r*) occurred at *r* > 30 pixels (15*μ*m) was deemed to be misaligned by more than one cell diameter and discarded.

The second quality control metric was based on the Wasserstein distance between the fiducial markers, *Wass*_*Epi* 1,*Epi* 2_. Since the majority of ROIs had 0 ≤ *Wass*_*Epi* 1,*Epi* 2_ ≤ 150, and in line with manual alignment assessment, we used *Wass*_*Epi* 1,*Epi* 2_ = 200 as an upper bound and discarded any ROIs with Wasserstein distances between the fiducial markers greater than this value. Choice of threshold parameters for both quality control checks was based on manual assessment. After quality control, 12,231 ROIs were deemed well aligned.

Approximate tissue area within each ROI was estimated by calculating the alpha hull of the point cloud (Edelsbrunner, Kirkpatrick, and Seidel 1983) with *α* = 500, and any ROIs whose tissue area was less than 80% of the 1000 x 1000 pixel area were discarded, leaving a final cohort of 43 patients with collectively 10,027 (49.6%) ROIs used for statistical analysis.

Following alignment quality control 2 of the 43 patients contain ROIs only within carcinoma regions and were subsequently excluded from intra-patient statistical analysis.

Together, these complementary spatial quality control checks ensure that ROIs included for spatial analysis can be robustly analysed using all spatial metrics, without introducing edge-effects caused by missing tissue or adenoma/carcinoma annotation boundaries.

#### UMAP

Dimension reduction for visualisation via UMAP was conducted using UMAP (Uniform Manifold Approximation and Projection) via the python module `umap`, after rescaling using the scikit-learn (`sklearn`) (Pedregosa et al. 2011) module via the function `sklearn.preprocessing.StandardScaler()`.

#### Principal Component Analysis

Dimension reduction for visualisation via PCA was conducted using the python module scikit-learn (`sklearn`) via the function `sklearn.decomposition.PCA()`.

#### Random Forests

Random forest classifiers (RF) were generated using the python module scikit-learn (`sklearn`). Data was divided into test and train datasets using sklearn.model_selection.train_test_split with a test_size of 0.33, and random forests trained using `sklearn.ensemble.RandomForestClassifier()` with n_estimators=1000 and other parameters at default values. Accuracy scores for each classifier were reported based on their performance predicting classes in the withheld test dataset.

#### Feature Importance Analysis

Feature importance scores are reported according to the Mean Decrease in Impurity (MDI) for each feature, obtained via `rf.feature_importances_` for a given random forest classifier `rf`. MDI (also known as Gini importance) is a normalised measure of how much each feature in the RF contributes to increasing the “purity” of the data when it is used in a decision tree in the RF. Features which do a good job of separating the adenoma and carcinoma classes when used in a decision tree will decrease the impurity of the input data by allocating adenoma and carcinoma regions down different sides of the decision tree. For more information, see (Pedregosa et al. 2011).

Importance scores of cell-cell interactions in adenoma and carcinoma classification were determined by the accumulation of all spatial metric feature importance scores that are associated with that ordered cell pair. The importance scores corresponding to cell counts are included within the aggregated importance score of cell-cell interactions of the same type (noting that for cells of the same type, the Wasserstein distance is identically 0 and the QCM is undefined).

#### Statistical Analysis

Statistical differences between spatial feature importances in figures 4G-4L were conducted using the two-sided Mann-Whitney-Wilcoxon test. These tests were calculated using the `statannotations.Annotator()` function with test=’Mann-Whitney’ from the `statannotations` python library(Charlier, Florian et al. 2022). Throughout the study, the resultant p-values (p) of these tests were grouped as follows: ns if *p* > 0.05, * if 0.01 < *p* ≤ 0.05, ** if 0.001 < *p* ≤ 0.01, *** if 0.0001 < *p* ≤ 0.001 and **** if *p* ≤ 0.0001.

## Acknowledgments

Oxford Applied Spatial Biology Initiative: Francesco Barbara, Maria Jose Jimenez Rodriguez, Sergio Serrano De Haro Ivanez, George William Atkinson, Nicholas Fan, Nicholas Lai. JAB and JWM were supported by Cancer Research UK (CR-UK) grant number CTRQQR-2021\100002, through the Cancer Research UK Oxford Centre. EJM is funded by the Lee Placito Medical Research Fund (University of Oxford). BJS and HMB are members of the Centre for Topological Data Analysis and this research was funded in whole or in part by EPSRC EP/R018472/1. BJS was further supported by the L’Oréal-UNESCO UK and Ireland For Women in Science Rising Talent Programme. SJL was supported by CRUK Programme Grant (DRCNPG-Jun22\100002). Core funding to the Wellcome Centre for Human Genetics was provided by the Wellcome Trust (090532/Z/09/Z). HJ was supported by Bowel Disease Research Foundation-Research Grant. JB was supported by the Oxford Clarendon Fund and the Queen’s College Graduate Fellowship.

## Author contributions

JAB, EJM, HMB, and SJL conceived and designed the project. Funding was obtained by SJL and EJM. Conceptual input and data interpretation was provided by all authors. Tissue curation and annotation was conducted by HJ, AE, SJL, and CC. Data curation and analysis was conducted by JAB, EJM, JWM, JJB, BJS, HRE, and JB. The manuscript was written by JAB, EJM, JWM, HMB, and SJL, and reviewed by all authors.

